# Wax ester production in nitrogen-rich conditions by metabolically engineered *Acinetobacter baylyi* ADP1

**DOI:** 10.1101/735274

**Authors:** Jin Luo, Elena Efimova, Pauli Losoi, Ville Santala, Suvi Santala

## Abstract

Metabolic engineering can be used as a powerful tool to redirect cell resources towards product synthesis, also in conditions that are not optimal. An example of a synthesis pathway strongly dependent on external conditions is the production of storage lipids, which typically requires high carbon/nitrogen ratio. *Acinetobacter baylyi* ADP1 is known for its ability to produce industrially interesting storage lipids, namely wax esters (WEs). Here, we engineered the central carbon metabolism of *A. baylyi* ADP1 by deletion of the gene *aceA* encoding for isocitrate lyase in order to allow redirection of carbon towards WEs. The production was further enhanced by overexpression of fatty acyl-CoA reductase Acr1 in the wax ester production pathway. This strategy led to 3-fold improvement in yield (0.075 g/g glucose) and 3.15-fold improvement in titer (1.82 g/L) and productivity (0.038 g/L/h) by a simple one-stage batch cultivation with glucose as carbon source. The engineered strain accumulated up to 27% WEs of cell dry weight. The titer and cellular WE content are the highest reported to date among microbes. We further showed that the engineering strategy alleviated the inherent requirement for high carbon/nitrogen ratio and demonstrated the production of wax esters using nitrogen-rich substrates including casamino acids, yeast extract and baker’s yeast hydrolysate, which support biomass production but not WE production in wild-type cells. The study demonstrates the power of metabolic engineering in overcoming natural limitations in the production of storage lipids.

## 1. Introduction

Metabolic engineering provides a powerful tool for improving the production of industrially relevant products using microbial cells as factories (Pfleger et al., 2015). One potential approach is to optimize the distribution of carbon and cellular resources between biomass and product synthesis, which is a common challenge in microbial production. Microbial storage lipid production represents a typical example of a synthesis route which strongly competes with biomass production. To induce lipid accumulation, excess of carbon along with the limitation of some key nutrients, typically nitrogen, is required (Breuer et al., 2012; Garay et al., 2014; Wältermann and Steinbüchel, 2005). Therefore, the production process is typically divided into two phases, where biomass is accumulated in the first phase and the lipid accumulation is induced in the second phase by limiting nitrogen availability while sustaining high carbon substrate concentration. However, such approach can result in low overall productivity. In addition, nitrogen limitation may not be suitable for cells (over)expressing non-native activities and pathways. One strategy to manipulate the distribution of carbon and energy between biomass and product synthesis is to directly engineer the central carbon metabolism. However, deletion of key genes in central metabolism may harm cell functions and strongly reduce growth, which can negatively affect production. Thus, in previous studies, various strategies for dynamic control of enzyme levels in central metabolism have been introduced. For example, Soma et al. increased the availability of acetyl-CoA from the tricarboxylic acid (TCA) cycle for isopropanol production by conditionally switching off the expression of citrate synthase (Soma et al., 2014).

In another example, Doong et al. improved the production of glucaric acid by dynamically down-regulating an essential enzyme in the competing glycolytic pathway based on quorum sensing, and up-regulating a downstream enzyme upon the production of its intermediate (Doong et al., 2018). More recently, Santala et al. established autonomous downregulation of the glyoxylate cycle based on the gradual oxidation of the inducer arabinose, enabling a shift from biomass formation to lipid accumulation (Santala et al., 2018).

Wax esters (WEs) are an example of high-value storage lipids, which could be used in a broad range of applications, including cosmetics, lubricants, pharmaceuticals, printing, food industries, etc. (Sánchez et al., 2016; Singh et al., 2007). *Acinetobacter* sp. is the best known microorganism for producing long-chain WEs as energy and storage compounds (Fixter et al., 1986; Rontani, 2010). In addition, *Acinetobacter baylyi* ADP1, is an ideal cellular platform for synthetic biology and metabolic engineering due to its genetic tractability (de Berardinis et al., 2008; Metzgar, 2004; Murin et al., 2012; Suárez et al., 2017; Tumen-Velasquez et al., 2018), thus enabling it to be a suitable host to study storage lipid production (Santala et al., 2014). Besides WEs, *A. baylyi* ADP1 has also been engineered to produce various native and non-native oleochemicals, such as alkanes, alkenes, and triacylglycerols (Lehtinen et al., 2018b; Luo et al., 2019; Salmela et al., 2019; Santala et al., 2011b).

Fatty acid synthesis pathway provides the precursors fatty acyl-CoAs (coenzyme A) for the synthesis of WEs (Janßen and Steinbüchel, 2014). The pathway produces fatty acyl-ACPs (acyl carrier protein) as products by iterative enzymatic reaction cycles, starting with the central metabolic intermediate, acetyl-CoA. In *Acinetobacter* sp., the resulting fatty acyl-ACPs are further converted to fatty acyl-CoAs, followed by three enzymatic reactions involved in WE synthesis: (1) reduction of fatty acyl-CoAs to fatty aldehydes by the fatty acyl-CoA reductase Acr1 (Reiser and Somerville, 1997), (2) reduction of the fatty aldehydes to fatty alcohols by an uncharacterized enzyme and (3) esterification between the fatty alcohols and the fatty acyl-CoAs by a bifunctional wax ester synthase/diacylglycerol acyltransferase (WS/DGAT) (Kalscheuer and Steinbüchel, 2003). The glyoxylate cycle, which widely exists in bacteria, hypothetically competes with fatty acid synthesis pathway for acetyl-CoA (Figure 1A). Unlike the complete TCA cycle, glyoxylate cycle bypasses two oxidative steps that release two molecules of carbon dioxide (Kornberg, 1966). Thus, one important role of the glyoxylate cycle is to replenish the intermediates in the TCA cycle that are withdrawn for growth, using two molecules of acetyl-CoA as substrate (Kornberg, 1966). The glyoxylate cycle is essential for cells growing on acetate or other non-glycolytic carbon sources, such as fatty acids, that are directly catabolized into acetyl-CoA. However, when glucose or amino acids are available, the glyoxylate cycle is not essential since the compounds required for growth can be supplied by alternative ways. In these cases, blocking the glyoxylate cycle can be hypothesized to enhance lipid production by increasing the flow of the available acetyl-CoA to fatty acid synthesis pathway and energy generation without affecting cell growth (Figure 1A).

**Figure 1.**
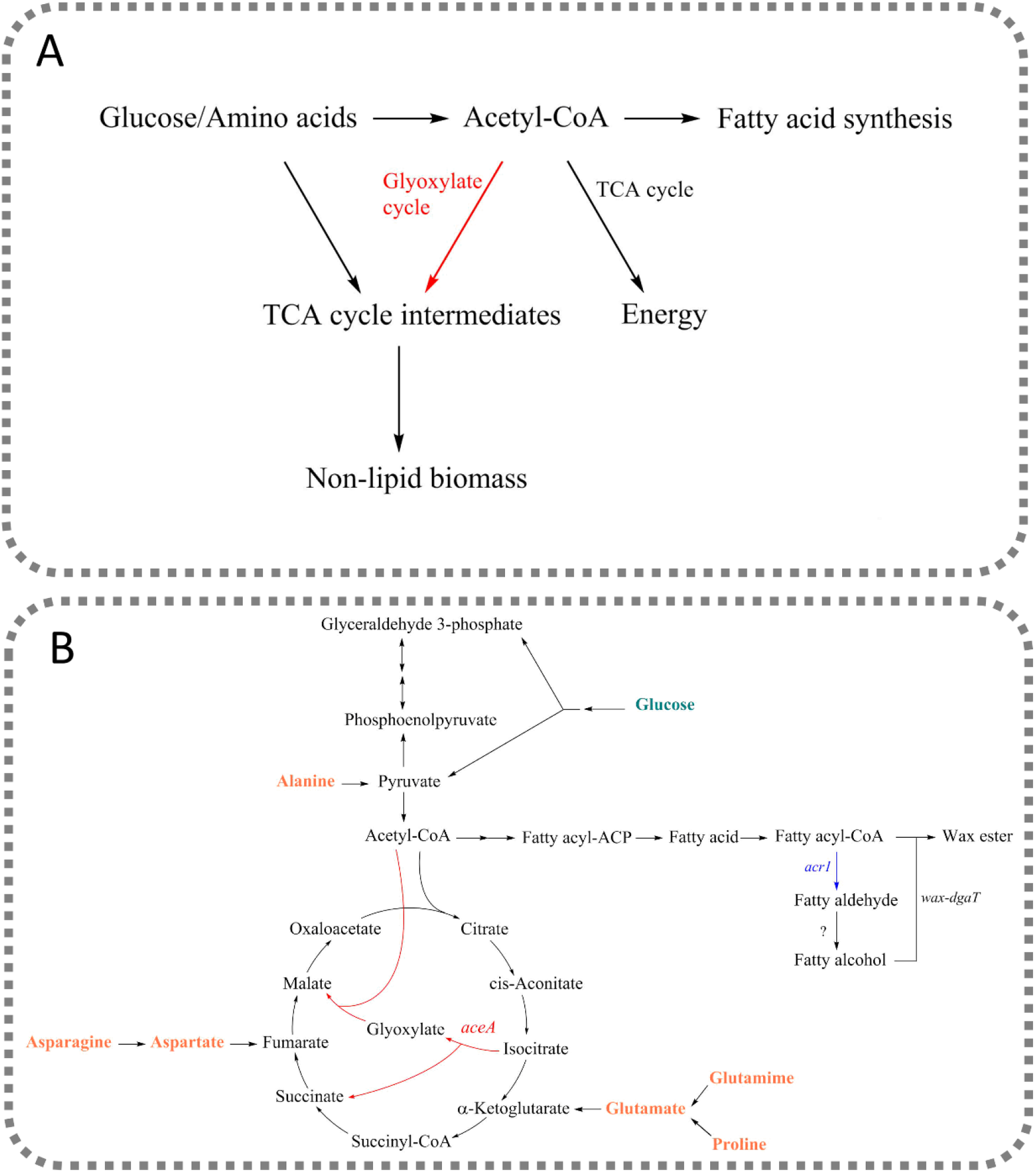
(A) Removal of the glyoxylate cycle hypothetically redirects the central metabolite, acetyl-CoA, from non-lipid biomass formation to fatty acid synthesis and energy generation. (B) Metabolic pathway for the synthesis of WEs from glucose and amino acids by *A. baylyi* ADP1. In the engineered strain, the gene *aceA* encoding for the isocitrate lyase, was deleted, blocking the glyoxylate cycle. The gene *acr1*, encoding for the fatty acyl-CoA reductase, was overexpressed to facilitate WE production.

For ADP1, protein-rich substrates provide an excellent carbon and nutrient source for growth, i.e. biomass production. However, due to the high nitrogen content, they are not suitable for the production of WEs. Here, we employ a metabolic engineering strategy aiming at efficient lipid accumulation, even from nitrogen-rich carbon sources, thus bypassing the need for high carbon/nitrogen ratio. We engineered *A. baylyi* ADP1 by deleting the gene *aceA* from the genome and overexpressing the gene *acr1* which encodes for the fatty acyl-CoA reductase (Figure 1B). The gene *aceA* encodes for isocitrate lyase, catalyzing the first step of the glyoxylate cycle, in which isocitrate is cleaved into succinate and glyoxylate. Glyoxylate is subsequently combined with one molecule of acetyl-CoA to form malate. The downstream pathway was up-regulated by *acr1* overexpression for directing the generated acetyl-CoA to WEs. We first evaluated the production of WEs by the engineered strain using glucose as carbon source. Prior to the evaluation of WE production from nitrogen-rich substrates, we examined the utilization of 20 amino acids as sole carbon source and explored the effect of *aceA* deletion on cell growth and lipid synthesis. We finally cultivated the engineered strain with different nitrogen-rich substrates (casamino acids, yeast extract and baker’s yeast hydrolysate) for WE production. We demonstrate that our approach significantly improved the production of WEs from both glucose and nitrogen-rich substrates.

## 2. Materials and methods

### 2.1. Strains and media

*Acinetobacter baylyi* ADP1 (DSM 24193, DSMZ, Germany) was used in the study for WE production. *E. coli* XL1-Blue (Stratagene, USA) was used as a host in plasmid constructions and amplifications. All strains used in the study are listed in Table 1.

**Table 1.**
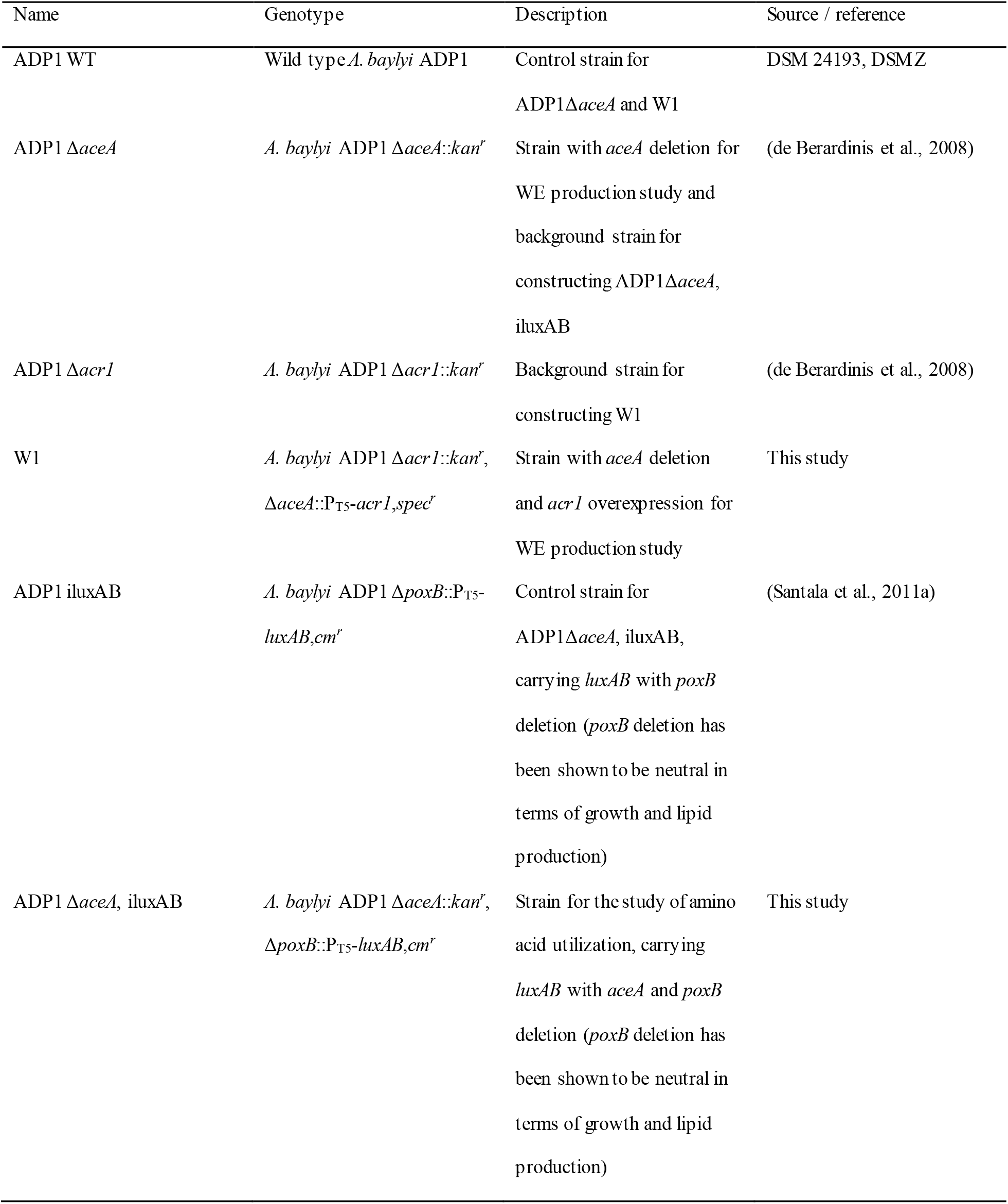
Bacterial strains used in the study

Modified LB medium (10 g/l tryptone, 5 g/l yeast extract, 1 g/l NaCl) supplemented with 1% glucose was used to grow *E. coli* and *A. baylyi* ADP1 for strain constructions. The cultivations for characterization of amino acid utilization and WE production were carried out using modified minimal salts medium MA/9 (Na_2_HPO_4_ 4.40 g/l, KH_2_PO_4_ 3.40 g/l, NH_4_Cl 1.00 g/l, nitrilotriacetic acid 0.008 g/l, NaCl 1.00 g/l, MgSO_4_ 240.70 mg/l, CaCl_2_ 11.10 mg/l, FeCl3 0.50 mg/l) supplemented with different types of carbon sources as appropriate. For the characterization of amino acid utilization, 20 L-amino acids (alanine, arginine, asparagine, aspartic acid, cysteine, glutamic acid, glutamine, glycine, histidine, isoleucine, leucine, lysine, methionine, phenylalanine, proline, serine, threonine, tryptophan, tyrosine, valine, all from Sigma) were added as sole carbon source respectively, with a concentration of 15 mM (except for tyrosine for which the concentration was made to 5 mM due to its low solubility). For WE production from glucose, 200 mM glucose and 2 g/L casamino acids were added. For WE production from nitrogen-rich substrates, casamino acids (10 g/L and 20 g/L), yeast extract (20 g/L), and 50% of baker’s yeast hydrolysate (the baker’s yeast was purchased from local market) were added, respectively. The pretreatment of the baker’s yeast was performed using the method described by Huo et al. (Huo et al., 2011), with some modifications. Briefly, 126 g/L biomass was mixed with water and heated at 80-85 °C for 18 min, after which the biomass was digested with 1-2 g/L protease (from *Bacillus* sp., Sigma) and incubated at 60 °C overnight. The digestion product was centrifuged, and the supernatant was collected and further filtered for use as substrate. Corresponding antibiotics were added in the media when necessary (with a concentration of 50 mg/L for spectinomycin, 50 mg/L for kanamycin, and 25 mg/L for chloramphenicol).

### 2.2. Genetic manipulation

The molecular cloning was performed using established methods. The reagents for the molecular work, including PCR, digestion and ligation, were provided by Thermo Scientific (USA) and used according to provider’s instruction. Transformation and genome editing of *A. baylyi* ADP1 were carried out as described previously (Santala et al., 2011b). All the primers used in the study are listed in Table S3.

The strains ADP1 Δ*aceA* (ΔACIAD1084) and ADP1 Δ*acr1* (ΔACIAD3383) were a kind gift from Dr Veronique de Berardinis (Genoscope, France). The strain ADP1 iluxAB was previously constructed in our laboratory (Santala et al., 2011a); in the strain the gene *poxB* (ACIAD3381) is replaced with the genes *luxAB* under T5 promoter. Deletion of the gene *poxB* has been shown to be neutral in terms of growth and WE production (Santala et al., 2011b).

The construction of the strain W1 was carried out as follows. First, the gene *acr1* was amplified from the genome of *A. baylyi* ADP1 using the primers tl17 and sa16. The amplified gene *acr1* was cloned to a previously described chromosomal integration cassette (Santala et al., 2011a) under a strong constitutive promoter T5. The original chloramphenicol resistance marker in the cassette was replaced with spectinomycin resistance marker originated from the plasmid pIM1463 (Murin et al., 2012), amplified with the primers tl45 and tl46. The resulting cassette was further amplified using the primers JL18-3 and JL18-4. The plasmid pUC57 containing the flanking sequences of the gene *aceA* was purchased from GenScript (USA) and further linearized using the primers JL18-1 and JL18-2. The linearized plasmid was ligated with the previously amplified cassette using Gibson Assembly, resulting in the targeting vector. The targeting vector was used transform the strain ADP1 Δ*acr1* to replace the gene *aceA* with the gene *acr1* under T5 promoter by homologous recombination, resulting the strain W1 (Δ*acr1:kan^r^*, Δ*aceA*::P_T5_-*acr1,spec^r^*). For the construction of the strain ADP1 Δ*aceA*, iluxAB, a previously constructed cassette (Santala et al., 2011a), which contains the genes *luxA* and *luxB* under T5 promoter, chloramphenicol resistance marker and the flanking sequences of the gene *poxB*, was used transform the strain ADP1 Δ*aceA*. The desired strains were selected with corresponding antibiotics.

### 2.3. Flux balance analysis

Flux balance analysis (FBA) (Orth et al., 2010) was used to predict the essentiality of gene deletion and calculate the theoretical yields of the conversion of the 6 L-amino acids (alanine, asparagine, aspartic acid, glutamic acid, glutamine and proline) and glucose into WEs by ADP1. A previously described genome-wide *A. baylyi* metabolic network (Durot et al., 2008) was used for the implementation of FBA. To allow simulation of WE accumulation, an exchange reaction of WEs was added in the metabolic network. In all conducted FBAs for essentiality prediction and WE yield calculation, the model was given 1 g/h/g CDW of each carbon source as an input (with the default minimal medium components and unlimited oxygen), and the growth reaction and the WE exchange reaction were maximized for respectively.

### 2.4. Characterization of amino acid utilization

The strains ADP1 Δ*aceA*, iluxAB and ADP1 iluxAB were used for the characterization of amino acid utilization. Both strains were cultivated in duplicate on 96-well plate (200 μl medium/well) with different single amino acids as carbon source respectively. The plate was incubated in Spark multimode microplate reader (Tecan, Switzerland) at 30 °C. Double orbital shaking was performed for 5 min twice an hour with an amplitude of 6 mm and a frequency of 54 rpm. Optical density at 600 nm (OD_600_) and luminescence signal were measured every 30 min.

### 2.5. Cultivation for wax ester production

For all the cultivations, the cells were first taken from agar plates containing LB medium and 1% glucose with corresponding antibiotics (if necessary, antibiotics were not added in subsequent cultivations). Cultivations were conducted in 50 ml medium/250 ml Erlenmeyer flasks at 25 °C and 300 rpm. Cultivations were carried out in duplicate with a starting OD_600_ of 0.1, using both glucose medium and nitrogen-rich media (media containing casamino acids, yeast extract and baker’s yeast hydrolysate respectively) as described in “Strains and media” section. For the cultivation with glucose, cells were cultivated for 24 h and 48 h before harvesting. Considering that WEs might be degraded after the carbon source is depleted, cells were harvested before reaching stationary phase in nitrogen-rich media which contain 10 g/L, 20 g/L casamino acids and 50% baker’s yeast hydrolysate. When cultivated with yeast extract, cells were harvested at 5 h (before reaching stationary phase) and 24 h.

### 2.6. Analytical methods

The baker’s yeast hydrolysate was prepared as described in “Strains and media” section. Protease was added for the hydrolysis of proteins into amino acids. The concentration of amine groups before and after protease digestion was determined by Ninhydrin Assay. To determine the protein content of the yeast sample, the biomass was heated in 0.5 N NaOH at 80-85 °C for 30 min, allowing release of protein. Bradford Assay was performed for protein concentration measurement.

WEs were analyzed by the thin layer chromatography (TLC) analysis after small-scale lipid extraction, as described earlier (Lehtinen et al., 2018a). Briefly, cells were collected from proper amount of culture by centrifugation. WEs were extracted from the cells with methanol-chloroform extraction method. The lower phase was used for TLC analysis, using glass HPTLC Silica Gel 60 F_254_ plate (Merck, USA). The mobile phase was a mixture of hexane, diethyl ether and acetic acid with a ratio of 90:15:1. Jojoba oil was used as the standard of WEs. Visualization was carried out with iodine.

Consumption of glucose was analyzed by high performance liquid chromatography (HPLC). The analysis was performed with LC-20AC prominence liquid chromatograph (Shimadzu, USA) equipped with RID-10A refractive Index detector. Phenomenex Rezex RHM-monosaccharide H+ (8%) column (Phenomenex, USA) was used and 0.01 N sulfuric acid was used as mobile phase with a pumping rate of 0.6 ml/min.

Quantification of WEs was carried out using ^1^H nuclear magnetic resonance (NMR). The cells were harvested from 40 ml culture by centrifugation at 30000 g for 30 min. The harvested cells were freeze-dried with Christ APLHA1-4 LD plus freeze dryer (Germany) for 24 h. The extraction of lipid from the freeze-dried cells and analysis with NMR were performed as described previously (Santala et al., 2011a). The chemical shift of 4.05 ppm corresponds to the proton signal of α-alkoxy-methylene group of WEs. The integral value of the signal at 4.05 ppm was used to calculate the content of WEs. An average molar mass of 506 g/mol of WEs was used to calculate the titer, yield, productivity, and content (Lehtinen et al., 2018a).

## 3. Results and discussion

### 3.1. Metabolic engineering for redirecting carbon flow from growth to lipid synthesis

Commercialization of microbial lipid production requires development of efficient cell catalyst. Previous metabolic engineering strategies have successfully demonstrated optimizing the production organisms, such as by blocking the competing pathways (Lu et al., 2008), increasing the supply of precursors (Lu et al., 2008; Tai and Stephanopoulos, 2013), deregulation of fatty acid synthesis pathway (Lu et al., 2008; Zhang et al., 2012) or balancing the reducing power (Qiao et al., 2017). However, challenges related to the competition between growth and lipid synthesis still remain. In addition, the tendency to accumulate lipids in oleaginous species, such as oleaginous yeast, depends on nitrogen limitation due to strict regulation (Garay et al., 2014). Typically, nitrogen limitation coupled with excess of carbon is used for lipid accumulation. In the strategy, biomass is accumulated in the first stage and lipid accumulation occurs upon nitrogen exhaustion. On one hand, the two-stage process can lower the productivity, and nitrogen limitation may not be suitable for cells (over)expressing non-native activities. On the other hand, lipid production from nitrogen-rich substrates, which represent abundant by-products in current industrial scenarios (Li et al., 2018), can be difficult.

WEs have been previously produced by *A. baylyi* ADP1 from various carbon sources, including alkanes (Ishige et al., 2002), glucose (Kannisto et al., 2017; Lehtinen et al., 2018a), acetate and butyrate (Salmela et al., 2018). However, WE production from nitrogen-rich substrates has not been described, possibly due to the requirement of nitrogen limitation to induce lipid accumulation. We have previously observed WE production during exponential phase in *A. baylyi* ADP1 (Santala et al., 2018, 2011a), indicating that nutrient limitation is not a strict requirement for lipid accumulation. We hypothesized that proper redirection of carbon flow from growth to lipid synthesis could enable enhanced WE accumulation and improved WE production from both the optimal substrate, namely glucose, and also from non-optimal substrates such as nitrogen-rich substrates. To validate the hypothesis, we constructed a strain W1 in which the gene *aceA* was replaced with the gene *acr1* under strong constitutive promoter. The native *acr1* was deleted in the strain. Deletion of *aceA* blocks the glyoxylate cycle, increasing the availability of acetyl-CoA for fatty acid synthesis pathway and energy generation. The gene *acr1* encodes for a NADPH-dependent fatty acyl-CoA reductase, which reduces fatty acyl-CoA to fatty aldehyde.

It was previously shown that overexpression of *acr1* improved WE production with a titer of 0.45 g/L (yield of 0.04 g/g glucose) (Lehtinen et al., 2018a). In the previous study, acr1 was located under lac-inducible promoter and highest WE production was obtained with the highest IPTG-concentration studied. Thus here we decided to use the strong T5-promoter. With this engineered strain, we studied WE production from both glucose and nitrogen-rich substrates, namely casamino acids, yeast extract and baker’s yeast hydrolysate.

The essentiality of *aceA* deletion to the cells when growing on acetate, glucose and 20 common L-amino acids was predicted using FBA by maximizing for the growth reaction before and after *aceA* deletion. In the simulations, 7 amino acids could be used as sole carbon source by *A. baylyi* ADP1, including alanine, arginine, asparagine, aspartate, glutamate, glutamine and proline (subsequent experimental validation showed that arginine cannot be used as sole carbon source). As expected, the gene *aceA* was predicted to be essential when acetate was used as sole carbon source; the theoretical specific growth rate did not change after *aceA* deletion when glucose or amino acids were used as carbon source (Table S1). We further inspected the change of the theoretical maximum yield of WEs from glucose and amino acids when increasing the carbon flux through isocitrate cleavage reaction. The inputs of different carbon sources (glucose and different amino acids) were set to 1 g/h/g CDW, and WE accumulation was maximized for. The maximum yield achievable by the model was decreased with the increase of flux through isocitrate cleavage reaction (Table S2). As the yields obtained with basic FBA consider only (genome-scale) stoichiometry but not kinetics or exact thermodynamics, the effect magnitude of *aceA* deletion on WE production is not accurately reflected in the simulation.

### 3.2. WE production from glucose

We first carried out batch-cultivation in flask with 200 mM glucose as carbon source to evaluate the production of WEs by the three strains, ADP1 WT (wild type ADP1 as control), ADP1 Δ*aceA* (with *aceA* deletion) and W1 (with *aceA* deletion and *acr1* overexpression). The three strains were cultivated for 24 h and 48 h. As shown in Figure 2, significantly improved WE production was observed in W1. After 24 h, both ADP1 Δ*aceA* and W1 had higher WE content and yield than ADP1 WT, and ADP1 Δ*aceA* had similar WE content as W1 (Figure 2B and 2C). However, ADP1 Δ*aceA* grew poorly, resulting in low CDW (Figure 2D) and low WE titer (Figure 2A). The strain seemed to stop growing before 24 h and did not further consume glucose (Figure S1). The reason for this is unknown. It is worth noting, however, that after 24 h, the pH of the culture dropped significantly below 4 for ADP1 Δ*aceA* while the pH was above 6.5 for W1 and ADP1 WT (data not shown), which indicated impaired metabolism in ADP1 Δ*aceA*. W1 had lower CDW than ADP1 WT (Figure 2D) but with 1.6-fold higher WE titer after 24 h (Figure 2A). After 48 h, W1 produced the highest WE titer of 1.82 g/L (3.15-fold higher than ADP1 WT) and the highest yield of 0.075 g/g glucose reported to date (3-fold higher than ADP1 WT) (Figure 2A and 2B), accumulating up to 0.27 g/g CDW of WEs (3-fold higher than ADP1 WT) (Figure 2C). The difference of CDW between the two strains was not significant (Figure 2D).

**Figure 2.**
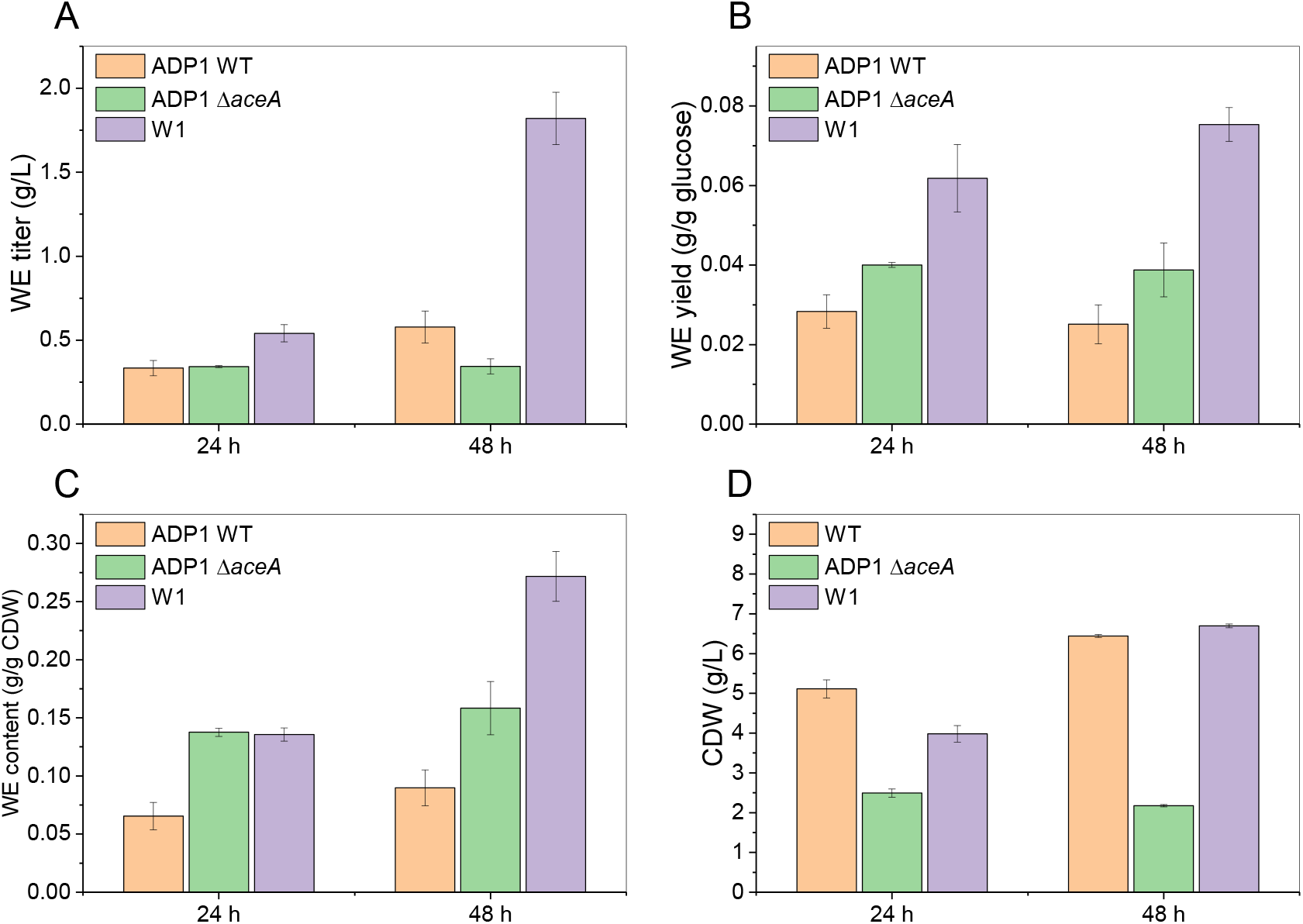
Comparison of (A) WE titer, (B) yield, (C) content and (D) CDW between the strains ADP1 WT, ADP1 Δ*aceA* and W1 after 24 h and 48 h of cultivation. The results represent the mean of two replicates and the error bars represent the standard deviations.

These results suggest that, when glucose was used as carbon source, *aceA* deletion in combination with *acr1* overexpression significantly enhanced WE accumulation without considerably reducing cell growth, resulting in improved titer, productivity and yield. Although the strain ADP1 *ΔaceA* grew poorly, it still had higher WE content and yield than ADP1 WT. However, it is not clear why ADP1 *ΔaceA* had poor growth and whether overexpression of *acr1* in W1 played a role in the rescue of growth. It can be speculated that the balance between energy production and biomass formation was impaired in the strain ADP1 *ΔaceA*. It was reported that, in *E. coli*, addition of glucose results in shut-off of the glyoxylate cycle due to the dephosphorylation of isocitrate dehydrogenase which competes with isocitrate lyase for the common substrate isocitrate (Walsh and Koshland, 1985). In this study, the altered phenotype of the *aceA* deletion strains when cultivated with glucose might indicate that in *A. baylyi* ADP1 the glyoxylate cycle is still active in the presence of glucose. If this is the case, the improved WE accumulation could be due to the increased availability of acetyl-CoA resulted from the blockage of the glyoxyate cycle as hypothesized. In addition, it is possible that some acetyl-CoA was redirected to the oxidative step in the TCA cycle, which might also benefit WE production by providing the reducing agent NADPH required for fatty acyl-ACP synthesis and fatty acyl-CoA reduction; *Acinetobacter sp*. has two isoenzymes of NADP+-dependent isocitrate dehydrogenase catalyzing the oxidation of isocitrate in the TCA cycle (Self and Weitzman, 1972).

The TCA cycle plays an important role in amino acid synthesis (Roberts et al., 1953). In *A. baylyi* ADP1, some TCA cycle intermediates, such as oxaloacetate and 2-ketoglutarate, are required for the synthesis of amino acids, such as aspartate, asparagine, glutamate, glutamine and proline. When glucose is used as carbon source, the intermediates of the TCA cycle are continuously withdrawn for amino acid synthesis and have to be replenished again. If the glyoxylate cycle is still active in the presence of glucose as suggested before, deletion of *aceA* will block the replenishment by the glyoxylate cycle from acetyl-CoA. As *A. baylyi* ADP1 lacks pyruvate carboxylase, the replenishment can be only accomplished by phosphoenolpyruvate carboxylation, in which phosphoenolpyruvate reacts with carbon dioxide to form oxaloacetate.

### 3.3. Exploration of amino acid utilization by the *aceA* knockout strain

Previous results indicate that *aceA* deletion increases the content and yield of WEs when glucose is used as carbon source. The same strategy is expected to benefit WE accumulation when cells are cultivated with nitrogen-rich substrates. Prior to using nitrogen-rich substrates for WE production, we first characterized the utilization of 20 single L-amino acids by the *aceA* knockout ADP1 and employed a bacterial luciferase-based biosensor to monitor the intracellular production of long-chain fatty aldehyde in real-time during the cultivation (Santala et al., 2011a). Bacterial luciferase (LuxAB) catalyzes the oxidation of FMNH_2_ (reduced flavin mononucleotide) and long-chain fatty aldehyde, resulting in emission of light (Szittner and Meighen, 1990). We integrated the gene *luxA* and *luxB* into the genome of the strain ADP1 Δ*aceA*, resulting in the strain ADP1 Δ*aceA*, iluxAB. A previously constructed strain ADP1 iluxAB was used as control (Santala et al., 2011a). Both strains were cultivated with 20 amino acids as sole carbon source respectively. Since long-chain fatty aldehyde is the precursor for WE synthesis, its production can reflect the carbon flux towards WE synthesis as well as fatty acid synthesis pathway.

The results showed that 6 amino acids could be used as sole carbon source by ADP1, including alanine, asparagine, aspartate, glutamate, glutamine and proline (Figure 3A). Arginine could not be used as sole carbon source by ADP1 although it was predicted to be used by the FBA (Table S1) due to the existence of the arginine succinyltransferase (AST) pathway in the ADP1 model.

**Figure 3.**
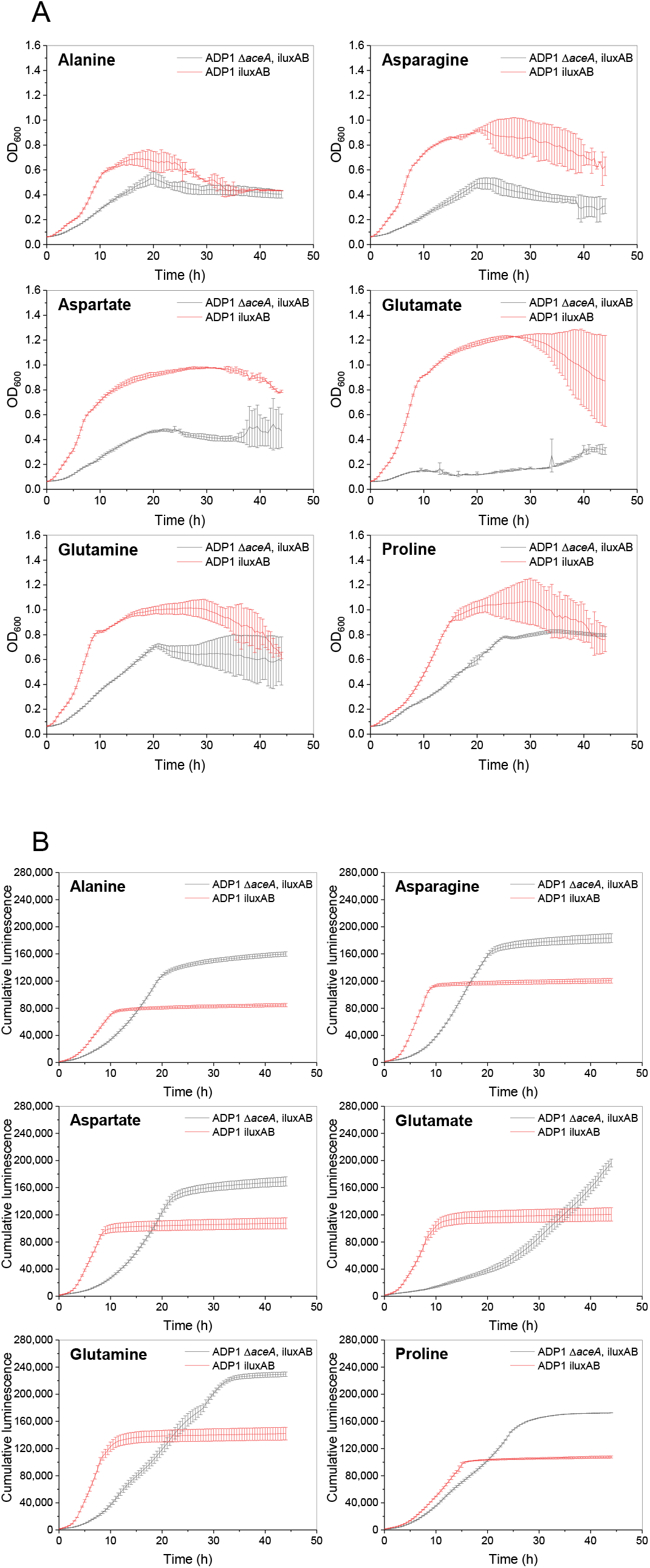
(A) Growth of the strains ADP1 iluxAB and ADP1 Δ*aceA*, iluxAB on alanine, asparagine, arsparate, glutamate, glutamine and proline respectively. The 6 amino acids can be used by *A. baylyi* ADP1 as sole carbon source. (B) Cumulative luminescence signal generated by ADP1 iluxAB and ADP1 Δ*aceA*, iluxAB when grown on the 6 amino acids respectively. No growth occurred with the other 14 individual amino acids. The results represent the mean of two replicates and the error bars represent the standard deviations.

In *E. coli*, the pathway allows arginine to be used as nitrogen source but not carbon source, putatively due to inappropriately controlled promoter or inadequate transport during carbon-limited growth (Schneider et al., 1998). It was previously hypothesized that the inability of ADP1 to use arginine as sole carbon source is due to the same reason (Durot et al., 2008). The 6 amino acids undergo different catabolic pathways. According to the metabolic model (Durot et al., 2008) used for FBA, L-alanine is first converted to D-alanine by alanine racemase, after which the D-alanine is converted to pyruvate by D-alanine dehydrogenase. Pyruvate can be converted to acetyl-CoA by pyruvate dehydrogenase complex or phosphoenolpyruvate by phosphoenolpyruvate synthase. In the *aceA* knockout strain, the acetyl-CoA cannot be converted to the TCA cycle intermediates through the glyoxylate cycle, and the replenishment is accomplished via phosphoenolpyruvate carboxylation, which is similar to the case with glucose. The catabolism of the other 5 amino acids leads to the formation of the TCA cycle intermediates; asparagine/aspartate is converted to fumarate or oxaloacetate while proline/glutamine/glutamate is converted to α-ketoglutarate. The TCA cycle intermediates malate oxaloacetate are converted to pyruvate and phosphoenolpyruvate by decarboxylation for gluconeogenesis and formation of acetyl-CoA.

As shown in Figure 3A, ADP1 Δ*aceA*, iluxAB grew more slowly and reached lower final OD_600_ than ADP1 iluxAB when cultivated with the 6 amino acids as sole carbon source. While the slower growth with alanine could be explained by lower availability of acetyl-CoA for the TCA cycle replenishment, it is surprising that the cultivations with the other 5 amino acids also led to slower growth since their catabolism forms the TCA cycle intermediates and does not directly lead to the formation of acetyl-CoA. Interestingly, the deletion of *aceA* seemed to have a negative effect on glutamate metabolism as the growth of ADP1 Δ*aceA*, iluxAB on glutamate was restrained to a much larger degree than on other amino acids (Figure 3A). In addition, diauxic growth was observed in ADP1 ΔaceA, iluxAB: the strain grew slowly in the beginning and stopped growing for a long period of time before growing again. The observation indicates that the deletion of *aceA* may have an influence on the regulation of glutamate metabolism.

The luminescence signals generated by the two strains were monitored during the cultivation; the cumulative luminescence can be used as an indicator to compare the amounts of fatty aldehyde produced by the two strains (Lehtinen et al., 2018a; Salmela et al., 2019; Santala et al., 2011a).

In comparison with ADP1 iluxAB, ADP1 Δ*aceA*, iluxAB generated higher cumulative luminescence in the end of the cultivations with not only alanine but also asparagine, aspartate, glutamine and proline (Figure 3B), suggesting more fatty aldehyde produced by ADP1 Δ*aceA*, iluxAB. This was also the case for the cultivation with glutamate, even though ADP1 Δ*aceA*, iluxAB grew poorly on glutamate. The change of luminescence/OD_600_ over time could enable us to elucidate the luminescence production profiles of the two strains (Figure S2). ADP1 iluxAB had similar luminescence production profiles on the six amino acids: the luminescence/OD_600_ peaked when cells were in early exponential phase, followed by drastic drop of the signal (Figure S2). It can be speculated that the cells were the most active in early exponential phase with available carbon flux towards fatty aldehyde, but the competition with growth for the carbon flux resulted in decrease of fatty aldehyde production. In comparison, the change of luminescence/OD_600_ was more subtle for ADP1 ΔaceA, iluxAB during cell growth (Figure S2), which might indicate more balanced distribution of carbon flux between growth and fatty aldehyde production. These results suggest that deletion of *aceA* could allow higher fatty aldehyde production at the cost of slower growth when single amino acids were used as sole carbon source.

### 3.4. WE production from casein amino acids and yeast extract

We next studied WE accumulation from casamino acids (casein hydrolysate) by the strain W1 and ADP1 WT. The strains W1 and ADP1 WT were cultivated in two conditions, with 10 g/L and 20 g/L casamino acids as carbon source. In order to compare WE accumulation between the two strains, the cells were harvested when OD_600_ was about 2 at which the cells were still in exponential phase, considering that WEs might be rapidly degraded under carbon limiting conditions (Santala et al., 2011a). As shown in Figure 4A, the concentration of casamino acids did not have a significant influence on the growth of both strains. ADP1 WT grew faster and reached OD_600_ of 2 after 4 h while W1 had a slower growth and reached OD_600_ of 2 after 11 h. As glutamic acid typically accounts for a large proportion in the amino acid profile of casein, it is possible that the negative effect on glutamate metabolism by *aceA* deletion may play a role in the slower growth of W1. Despite being in nitrogen-rich conditions, WT ADP1 still accumulated a small amount of WEs during exponential phase, with a content of 0.02 g/g CDW in both 10 and 20 g/L casamino acids (Figure 4B). In comparison, W1 accumulated approximately 0.13 and 0.11 g/g CDW of WEs in the cultivations with 10 g/L and 20 g/L casamino acids, respectively (Figure 4B), suggesting 5-6 folds enhanced accumulation of WEs during exponential phase. The contents were similar to that obtained with ADP1 WT from glucose.

**Figure 4.**
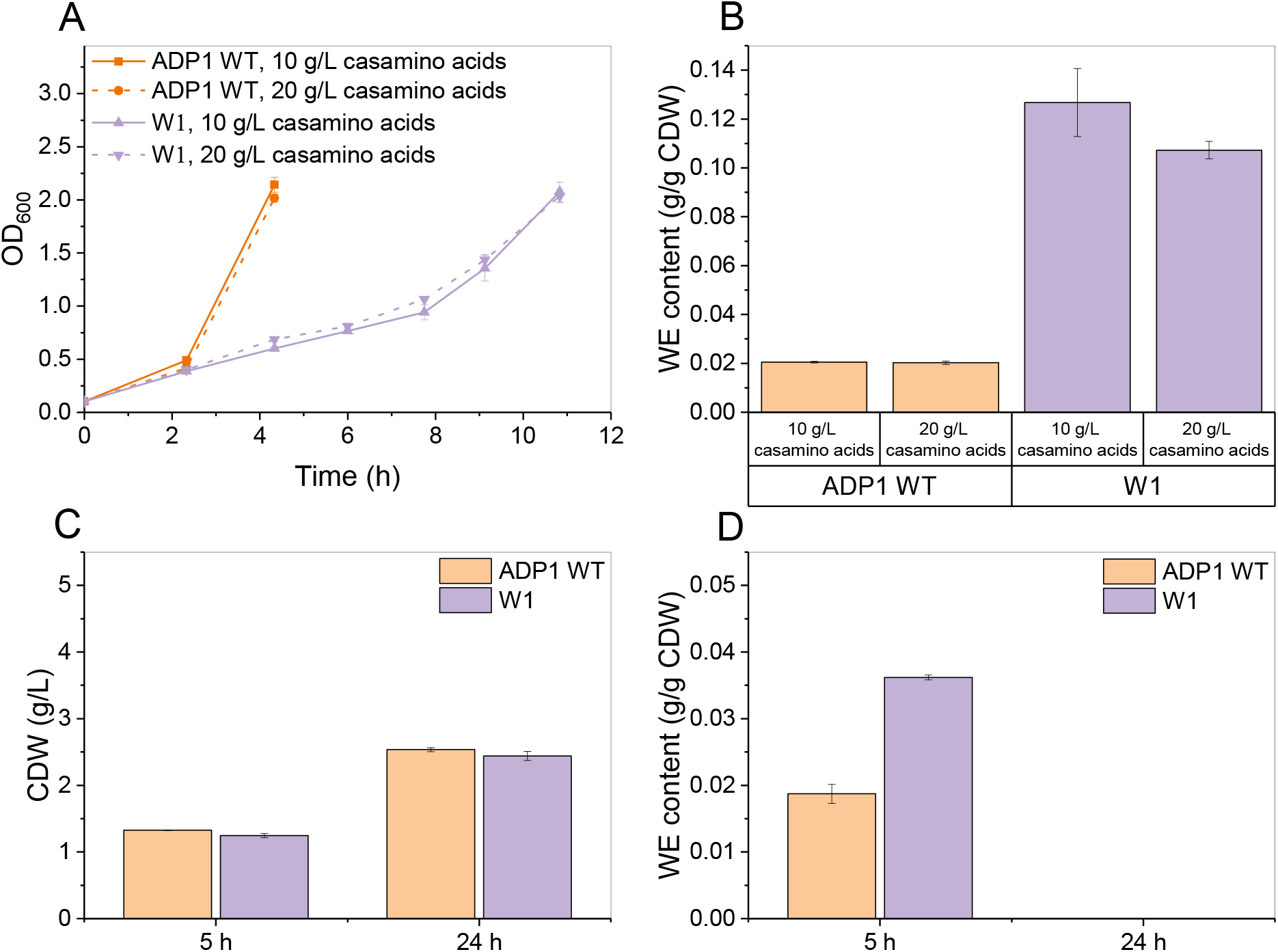
(A) Growth (represented as OD_600_) of the strains W1 and ADP1 WT in 10 and 20 g/L casamino acids respectively. (B) Comparison of WE content between W1 and ADP1 WT in exponential phase when grown in 10 and 20 g/L casamino acids. Cells of both strains were harvested when in exponential phase; ADP1 WT was harvested after 4 h, and WI was harvested after 11 h. Comparison of (C) CDW and (D) WE content between W1 and ADP1 when cultivated with 20 g/L yeast extract. The results represent the mean of two replicates and the error bars represent the standard deviations.

To study the WE accumulation using yeast extract, a cultivation with 20 g/L yeast extract as substrate was performed. The cells were harvested during exponential phase (5 h) and after reaching stationary phase (24 h). Interestingly, W1 and ADP1 WT showed similar growth, and the difference of CDW between the two strains was not apparent when harvested after 5 h of cultivation (Figure 4C). Although W1 had 1.9-fold higher WE content than ADP1 WT (Figure 4D), the content (0.036 g/g CDW) was much lower than those obtained with casamino acids. The difference of cell growth phase may affect the content of WEs, but the main reason of the lower WE content might be attributed to the composition difference between casamino acids and yeast extract. Apart from free amino acids and peptides, yeast extract also contains other nutrients, such as vitamins, macro and micro minerals, which may benefit growth more than lipid accumulation. After 24 h, WEs were not detected from both strains due to degradation (Figure 4D). Under the nitrogen-rich condition, carbon source was likely to be the growth-limiting factor during the stationary phase, and the accumulated WEs could be consumed as energy and carbon source. Further engineering, such as blocking β-oxidation by deletion of related genes (Lu et al., 2008; Steen et al., 2010), could be a potential strategy to prevent WE degradation.

Here, we demonstrated a novel metabolic engineering strategy that resulted in significantly improved WE production in nitrogen-rich conditions. However, further engineering is still needed to obtain high WE titer from protein-rich substrates. The *A. baylyi* strain employed in the study can only utilize 6 amino acids as sole carbon source. For the application point of view, it is important to develop a strain that can utilize more amino acids as carbon source. Chemical mutagenesis has been shown to be an effective way to improve amino acid utilization (Huo et al., 2011), and the obtained *E. coli* mutant was able to utilize 13 individual amino acids as sole carbon source, compared to 4 for the wild type strain. Moreover, microbes have evolved mechanisms that favor amino acid anabolism (Wernick and Liao, 2013). A driving force for deamination would allow more carbon skeletons to be released from amino acids for product formation, and the strategies for creating the driving force have been well demonstrated in *E. coli* by Huo et al. (Huo et al., 2011). It was shown that deamination could be facilitated by deletion of genes related to ammonium-assimilation pathway (Huo et al., 2011; Mikami et al., 2017). Furthermore, introduction of exogenous transamination-deamination cycles further drained amino acids that serve as nitrogen reservoir as the result of transamination

### 3.5. WE production from baker’s yeast hydrolysate

Renewable plant-derived biomass and algal biomass have been emerging as alternatives to fossil fuels for the production of fuels and chemicals (Peralta-Yahya et al., 2012; Pfleger et al., 2015; Ragauskas, 2006; Steen et al., 2010). However, this scenario is accompanied with accumulation of large amount of nitrogen-rich by-products (Li et al., 2018). It was estimated that 100 Mt/year of protein could be generated if 10% of transportation fuels were produced from biomass-derived sources (Tuck et al., 2012). In addition, considerable amount of protein-rich wastes are produced from brewery and food industries (Ferreira et al., 2010; Tuck et al., 2012). It is of great importance to exploit the value of these waste streams on the ground of circular economy. One option is to use them as animal feeds, but the limited market is not able to deal with the increasing amount of protein wastes (Huo et al., 2011). Thus, strategies have been brought out for the valorization of protein wastes, of which microbial production of valuable products directly from protein wastes is advantageous, as compared to other processes that require costly amino acid isolation steps (Li et al., 2018). Although production of a range of chemicals from protein sources has been proposed (Huo et al., 2011), it is challenging to efficiently produce oleochemicals, such as storage lipids, from nitrogen-rich protein wastes.

To evaluate of possibility of using industrial spent yeast for WE production, dry baker’s yeast was used as a representative feedstock. To obtain baker’s yeast hydrolysate that can be used as substrate, a heat treatment was performed for the biomass, followed by digestion with protease to break the peptide bonds of released proteins. The medium was supplemented with 50% baker’s yeast hydrolysate for cultivation. Cells were harvested during exponential phase (after 4.7 h) and early stationary phase (after 8 h) where the WE content is likely to be the highest. The CDWs of the strains W1 and ADP1 WT did not differ significantly after 4.7 and 8 h of cultivation (Figure 5C). W1 accumulated WEs from 0.025 g/g CDW in exponential phase to 0.030 g/g CDW in early stationary phase, while the WE content of ADP1 WT did not significantly increase in the end of the cultivation and was maintained at around 0.016 g/g CDW (Figure 5B). The difference of WE content between the two strains can be also seen through WE visualization (Figure 5D). The final WE titer of 0.066 g/L was obtained with WI, which was 2-fold higher than that obtained with ADP1 WT (Figure 5A). The results suggest that the protein-rich biomass can be a good feedstock for microbial production; the fermentable hydrolysate could be obtained by simple pretreatment process.

**Figure 5.**
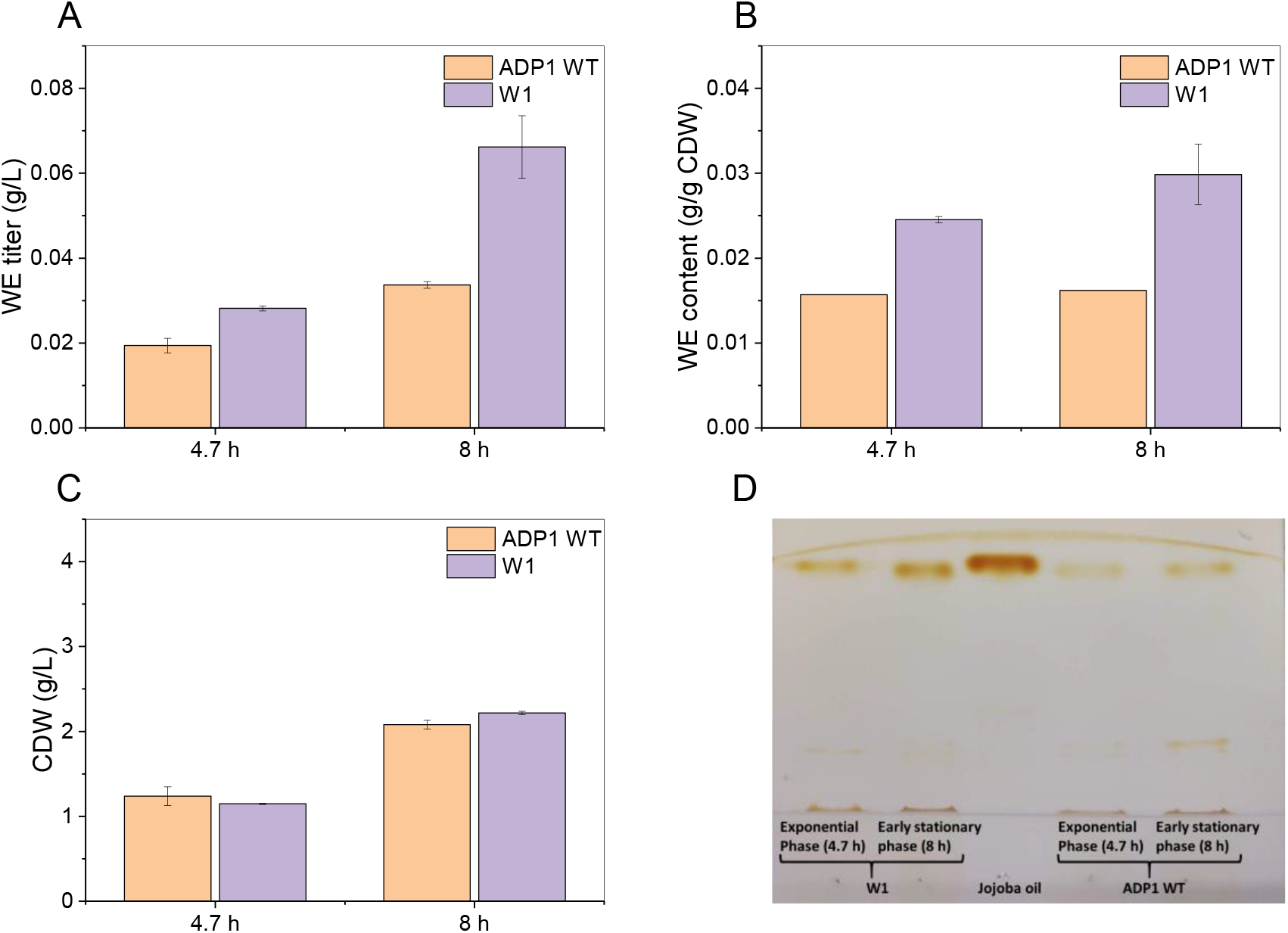
Comparison of (A) WE titer, (B) content and (C) CDW between ADP1 WT and W1 when grown in 50% baker’s yeast hydrolysate. The results represent the mean of two replicates and the error bars represent the standard deviations. (D) WE visualization by TLC (Thin-layer chromatography). Same amount of biomass was taken from each culture for lipid extraction. Jojoba oil was used as standard.

## 4. Conclusions

In this study, we engineered *A. baylyi* ADP1 to improve the production of WEs by deletion of *aceA* and overexpression of *acr1*. As a result of redirecting the carbon flux, the engineered strain produced 27% WEs of CDW with a titer of 1.8 g/l from glucose, representing approximately 3-fold improvement in production metrics (titer, yield, productivity) compared to WT. We further demonstrated the possibility to use nitrogen-rich substrates for WE production by *A. baylyi* ADP1. The engineering led to 5-6 fold improvement in WE accumulation from casamino acids; the amount of WEs produced by the engineered strain was similar to that produced by ADP1 WT from glucose. The study represents the starting point for establishing efficient high-value lipid production from non-optimal substrates such as protein-rich waste streams. We also anticipate that this approach could provide insight for improving the production of other oleochemicals.

## Author contribution

JL, SS, and VS designed the study. JL carried out the microbiological and molecular work. PL and JL conducted flux balance analysis. EE and JL conducted substrate and WE analysis. All authors participated in writing the manuscript. All authors read and approved the final manuscript.

## Abbreviations

WEs: wax esters
TCA: tricarboxylic acid
FBA: flux balance analysis

