## Supplementary materials for "Wax ester production in nitrogen-rich conditions by metabolically engineered *Acinetobacter baylyi* ADP1"

**Table S1.** Theoretical specific growth rate (1/h) of the strain ADP1 WT and ADP1 Δ*aceA* on different carbon sources. Flux balance analysis (FBA) was used for calculating the specific growth rate. The same genome-wide *A. baylyi* metabolic network was used for both strains in FBA, except that the gene *aceA* was deleted from the network for ADP1 Δ*aceA*. In all conducted FBAs, 1 g/h/g CDW of each carbon source was set as an input (with the default minimal medium components and unlimited oxygen), and the growth reaction was maximized for.

| | **ADP1 WT** | **ADP1 Δ*aceA*** |
|---|---|---|
| **Acetate** | 0.413 | 0 |
| **Ala** | 0.465 | 0.46 |
| **Arg** | 0.416 | 0.416 |
| **Asn** | 0.33 | 0.33 |
| **Asp** | 0.321 | 0.321 |
| **Cys** | 0 | 0 |
| **Glucose** | 0.453 | 0.453 |
| **Gln** | 0.451 | 0.451 |
| **Glu** | 0.438 | 0.438 |
| **Gly** | 0 | 0 |
| **His** | 0 | 0 |
| **Ile** | 0 | 0 |
| **Leu** | 0 | 0 |
| **Lys** | 0 | 0 |
| **Met** | 0 | 0 |
| **Phe** | 0 | 0 |
| **Pro** | 0.629 | 0.629 |
| **Ser** | 0 | 0 |
| **Thr** | 0 | 0 |
| **Trp** | 0 | 0 |
| **Tyr** | 0 | 0 |
| **Val** | 0 | 0 |

**Table S2.** Theoretical yield of WEs (expressed as g WEs/g carbon source) from different carbon sources by *A. baylyi* ADP1 (table on the left) and the theoretical yield when the flux through isocitrate cleavage reaction was increased from 0 mmol/h/g CDW to 5 mmol/h/g CDW (table on the right). FBA was used for calculating the theoretical yield. In all conducted FBAs, 1 g/h/g CDW of each carbon source was set as an input (with the default minimal medium components and unlimited oxygen), and the exchange reaction of WEs was maximized for.

| | |
|---|---|
| **Ala** | 0.305 |
| **Arg** | 0.17 |
| **Asn** | 0.21 |
| **Asp** | 0.204 |
| **Cys** | 0 |
| **Glucose** | 0.281 |
| **Gln** | 0.203 |
| **Glu** | 0.201 |
| **Gly** | 0 |
| **His** | 0 |
| **Ile** | 0 |
| **Leu** | 0 |
| **Lys** | 0 |
| **Met** | 0 |
| **Phe** | 0 |
| **Pro** | 0.257 |
| **Ser** | 0 |
| **Thr** | 0 |
| **Trp** | 0 |
| **Tyr** | 0 |
| **Val** | 0 |

| | Flux through isocitrate cleavage reaction ( mmol/h/g CDW) | | | | | | |
|---|---|---|---|---|---|---|---|
| | 0 | **0.1** | **0.25** | **0.5** | **1** | **2.5** | **5** |
| **Ala** | 0.305 | 0.305 | 0.305 | 0.304 | 0.303 | 0.258 | 0.184 |
| **Arg** | 0.17 | 0.167 | 0.163 | 0.155 | 0.14 | 0.096 | 0.022 |
| **Asn** | 0.21 | 0.21 | 0.21 | 0.209 | 0.195 | 0.15 | 0.076 |
| **Asp** | 0.204 | 0.204 | 0.204 | 0.203 | 0.193 | 0.149 | 0.074 |
| **Cys** | 0 | 0 | 0 | 0 | 0 | 0 | 0 |
| **Glucose** | 0.281 | 0.281 | 0.28 | 0.28 | 0.279 | 0.255 | 0.181 |
| **Gln** | 0.203 | 0.2 | 0.195 | 0.188 | 0.173 | 0.129 | 0.055 |
| **Glu** | 0.201 | 0.198 | 0.194 | 0.187 | 0.172 | 0.127 | 0.053 |
| **Gly** | 0 | 0 | 0 | 0 | 0 | 0 | 0 |
| **His** | 0 | 0 | 0 | 0 | 0 | 0 | 0 |
| **Ile** | 0 | 0 | 0 | 0 | 0 | 0 | 0 |
| **Leu** | 0 | 0 | 0 | 0 | 0 | 0 | 0 |
| **Lys** | 0 | 0 | 0 | 0 | 0 | 0 | 0 |
| **Met** | 0 | 0 | 0 | 0 | 0 | 0 | 0 |
| **Phe** | 0 | 0 | 0 | 0 | 0 | 0 | 0 |
| **Pro** | 0.257 | 0.254 | 0.25 | 0.243 | 0.228 | 0.183 | 0.109 |
| **Ser** | 0 | 0 | 0 | 0 | 0 | 0 | 0 |
| **Thr** | 0 | 0 | 0 | 0 | 0 | 0 | 0 |
| **Trp** | 0 | 0 | 0 | 0 | 0 | 0 | 0 |
| **Tyr** | 0 | 0 | 0 | 0 | 0 | 0 | 0 |
| **Val** | 0 | 0 | 0 | 0 | 0 | 0 | 0 |

**Table S3.** List of primers used in the study

| Name | Description | Oligo sequence (5-3') |
|---|---|---|
| tl17 | XbaI, amplification of the gene *acr1* (ACIAD3383) | TGGAATTCGCGGCCGCTTCTAGAGAAAGAGGAGAAATACTAGATGATATCAATCAGGGAAAAACGCG |
| sa16 | XhoI, amplification of the gene *acr1* (ACIAD3383) | CTTCTTCTCGAGTTATTACCAGTGTTCGCCTGG |
| tl45 | XhoI, amplification of spectinomycin resistance marker | GTAGCGCTCGAGGCAGAAAGGAGAAGCTTACTAGC |
| tl46 | PstI, amplification of spectinomycin resistance marker | ATCTTGCTGCAGCTCGGCTTGAACGAATTGTTAGAC |
| JL18_1 | Linearization of the plasmid pUC57 containing of the flanking sequences of the gene *aceA* for Gibson Assembly | GGCGTATGGTTTAAAAAAC |
| JL18_2 | Linearization of the plasmid pUC57 containing of the flanking sequences of the gene *aceA* for Gibson Assembly | GATATATTCCCTTTTAGGATTTC |
| JL18_3 | Amplification of the gene *acr1* (ACIAD3383) under T5 promoter for Gibson Assembly | ATCCTAAAAGGGAATATATCCAATTGGCTGGCATCCCTAAC |
| JL18_4 | Amplification of the gene *acr1* (ACIAD3383) under T5 promoter for Gibson Assembly | GGTTTTTTAAACCATACGCCCCTAGGGCTTAATGCGCC |
| 1084 V1 | Confirmation of the deletion of the gene *aceA* (ACIAD1084) | TTTTTCTATCATTCATTTTTAAGTC |
| 1084 V2 | Confirmation of the deletion of the gene *aceA* (ACIAD1084) | CTCAACATGATATGCACACTGC |

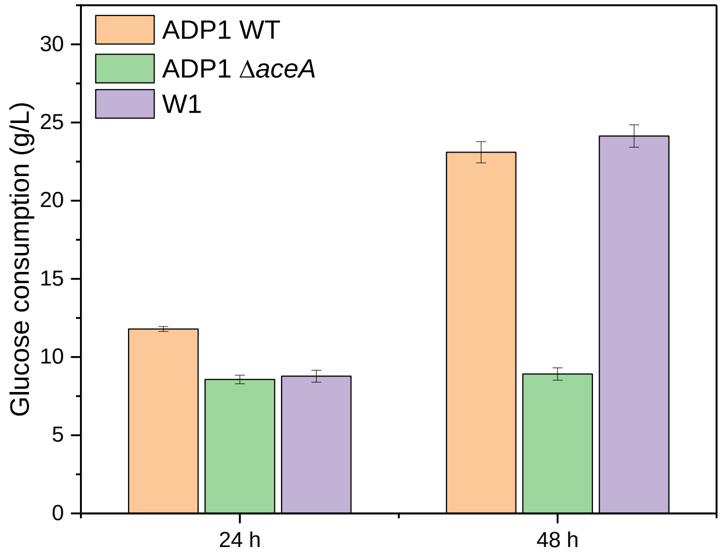


**Figure S1.** Consumption of glucose by ADP1 WT, ADP1 Δ*aceA* and W1 after 24, 48 and 72 h of cultivation. The initial concentration of glucose was 200 mM (corresponding to 36 g/L). Glucose was not completely consumed by all the three strains after 48 h of cultivation. The results represent the mean of two replicates and the error bars represent the standard deviations.

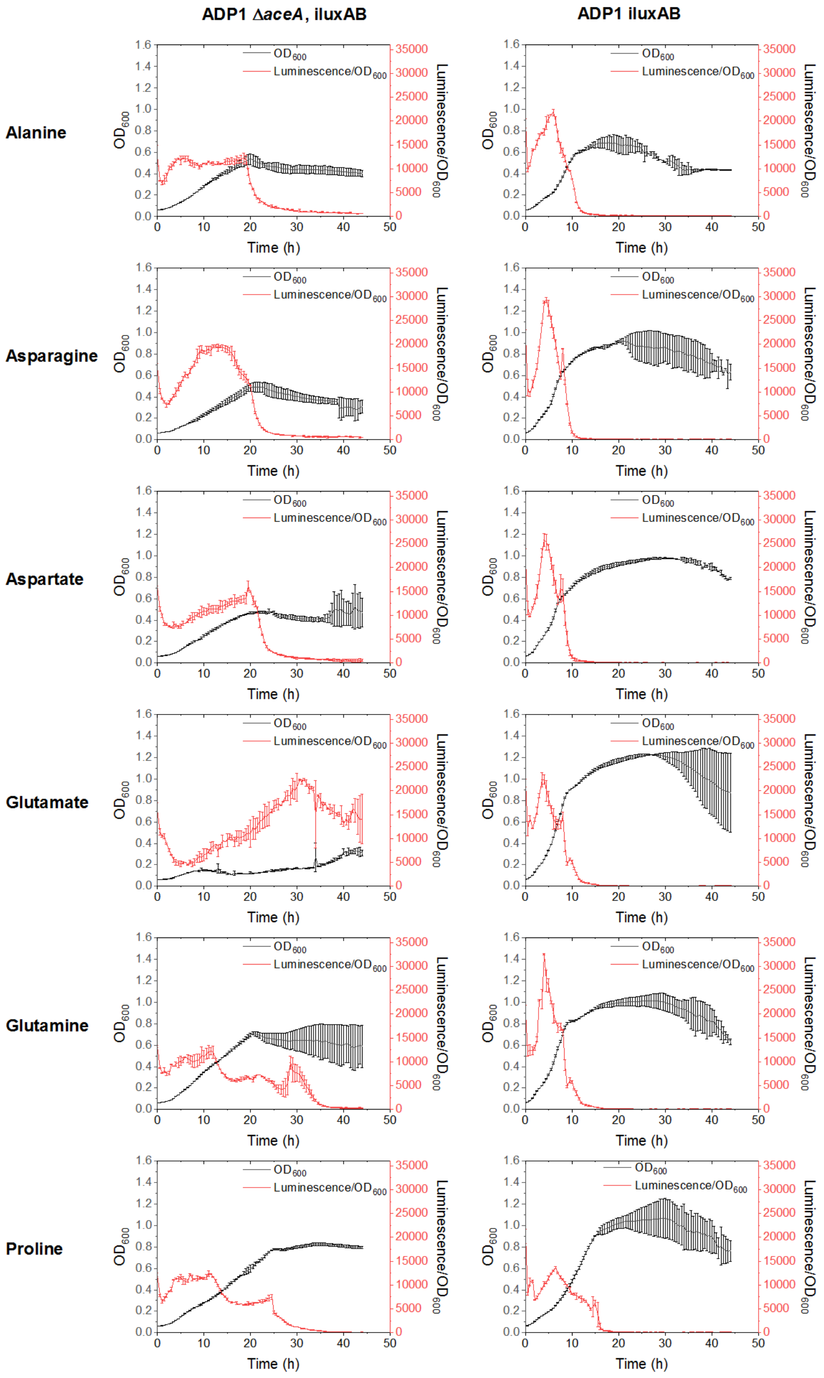


**Figure S2.** Change of luminescence/OD600 and OD600 over time for the strain ADP1 Δ*aceA*, iluxAB (first column) and ADP1 iluxAB (second column) on different amino acids (corresponding to different rows). Both strains were cultivated on 96-well plate with different single amino acids as carbon source respectively. OD600 and luminescence signal were measured every 30 min. The results represent the mean of two replicates and the error bars represent the standard deviations.

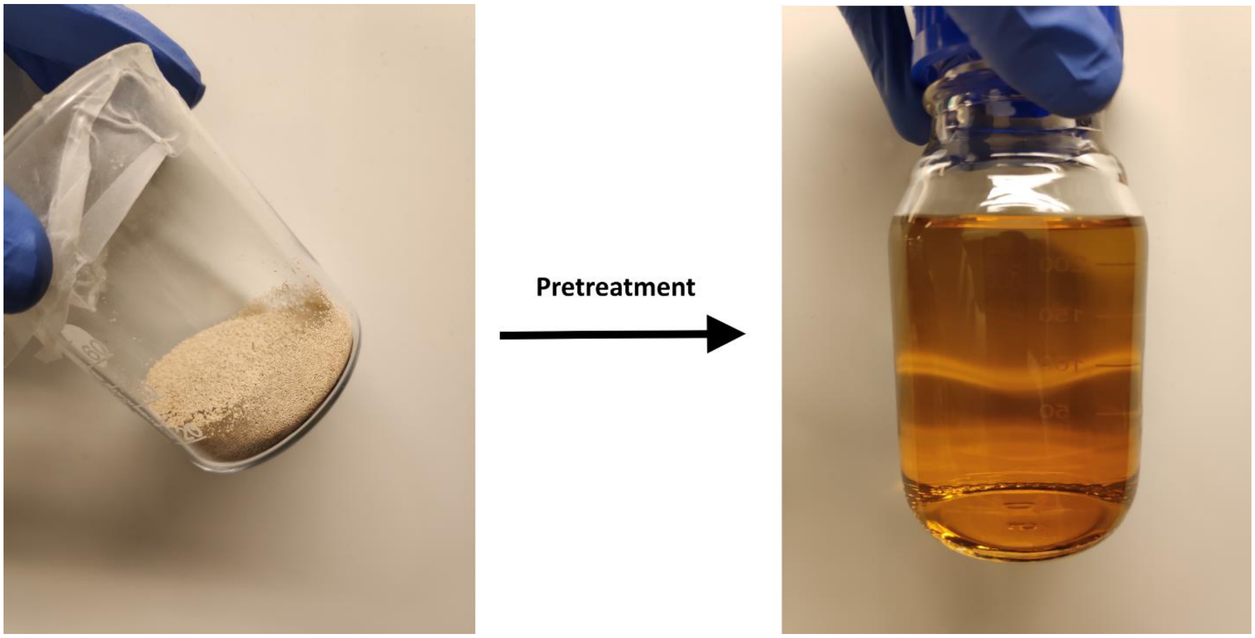


**Figure S3.** Dry baker yeast (on the left) and its hydrolysate stock solution (on the right). 126 g/L of the biomass was heated in water at 80-85 degree for 18 min, allowing release of 44582 µM free amine groups measured by Ninhydrin test. The biomass was then digested with 1-2 g/L g/L of protease at 60 degree overnight, further increasing the amount of amine group to 72450 µM. The hydrolysate was centrifuged, and the supernatant was filtered for use as substrate.
